# Fabrication of elastomeric stencils for patterned stem cell differentiation

**DOI:** 10.1101/2024.03.01.582929

**Authors:** Stefanie Lehr, Jack Merrin, Monika Kulig, Thomas Minchington, Anna Kicheva

**Author notes:** Equal contribution (Stefanie Lehr, Jack Merrin).

## Abstract

Stem cell differentiation with controlled geometry results in reproducible pattern formation. In contrast to constraining differentiating cells on micropatterned surfaces, we initialise colony formation using elastomeric stencils that adhere to culture dishes and create microwells with defined sizes and shapes. After colony formation, stencils are removed to allow colony growth and cell migration. Stencil fabrication involves mould production by photolithography followed by replica-moulding polydimethylsiloxane (PDMS). This approach produces reproducible two-dimensional organoids tailored for quantitative studies of growth control and pattern formation.

## Before you begin

### Background and motivation

The directed differentiation of embryonic stem (ES) cells into defined cell and tissue types has had a marked impact on tissue engineering as well as on studies of the principles of embryonic development and disease. In recent years, it has become increasingly clear that geometric constraints are essential to overcome the intrinsic heterogeneity of *in vitro* differentiation systems^1–3^. This principle has been applied to two-dimensional ES cell differentiation systems on restricted micropatterns to yield quantitative data on pattern formation and signalling, for instance, during mouse and human gastrulation^4–6^. In these systems, micropatterned cell culture surfaces are created by defining areas that promote cell attachment, passivated areas to prevent cell attachment, or some combination of the two^7,8^. While these systems provide a convenient quantitative readout of pattern formation, the confinement of cells within a restricted surface limits the applicability of this approach to investigating growing tissues and migratory cell populations.

We recently developed a method for directed differentiation of mouse ES cells into cell types of the developing dorsal spinal cord^9^. With this approach, cell types of the dorsal spinal cord form remarkable self-organised patterns upon exposure to BMP4. Trunk neural crest, a highly migratory cell type, forms at the periphery, followed by roof plate and dorsal neural progenitor subtypes dP1-6 towards the centre of colonies in their correct spatial order^9^. To allow for neural crest migration and reproducible two-dimensional patterning of the colonies, we use an approach that relies on removable PDMS microwell stencils^10,11^. This initialises colony formation on a defined area and subsequently allows colony growth and migration. Here, we provide an extended step-by-step protocol for stencil microfabrication adapted for the directed differentiation of mouse embryonic stem cells into dorsal spinal neural tube progenitors.

#### NOTE

Completing this protocol will require expertise in photolithography and standard approaches for working with PDMS for soft lithography. Typically, this involves the use of a cleanroom.

### Design and ordering of the photomask

#### NOTE

Photomasks are ordered from an external company; therefore, this step has to be completed in advance. We ordered photomasks from JDPhotoData (UK) as a 9 × 12” film photomask with a resolution of approximately 10 μm with positive polarity and emulsion film side down.

The photomask is used to fabricate a master silicon wafer mould for the stencils using photolithography. The transparent areas on the mask will become posts in the mould and, therefore, define the holes in the final stencils (Figure 1). Photomasks with the desired pattern can be designed using CAD software and ordered from a specialist supplier.

**Figure 1.**
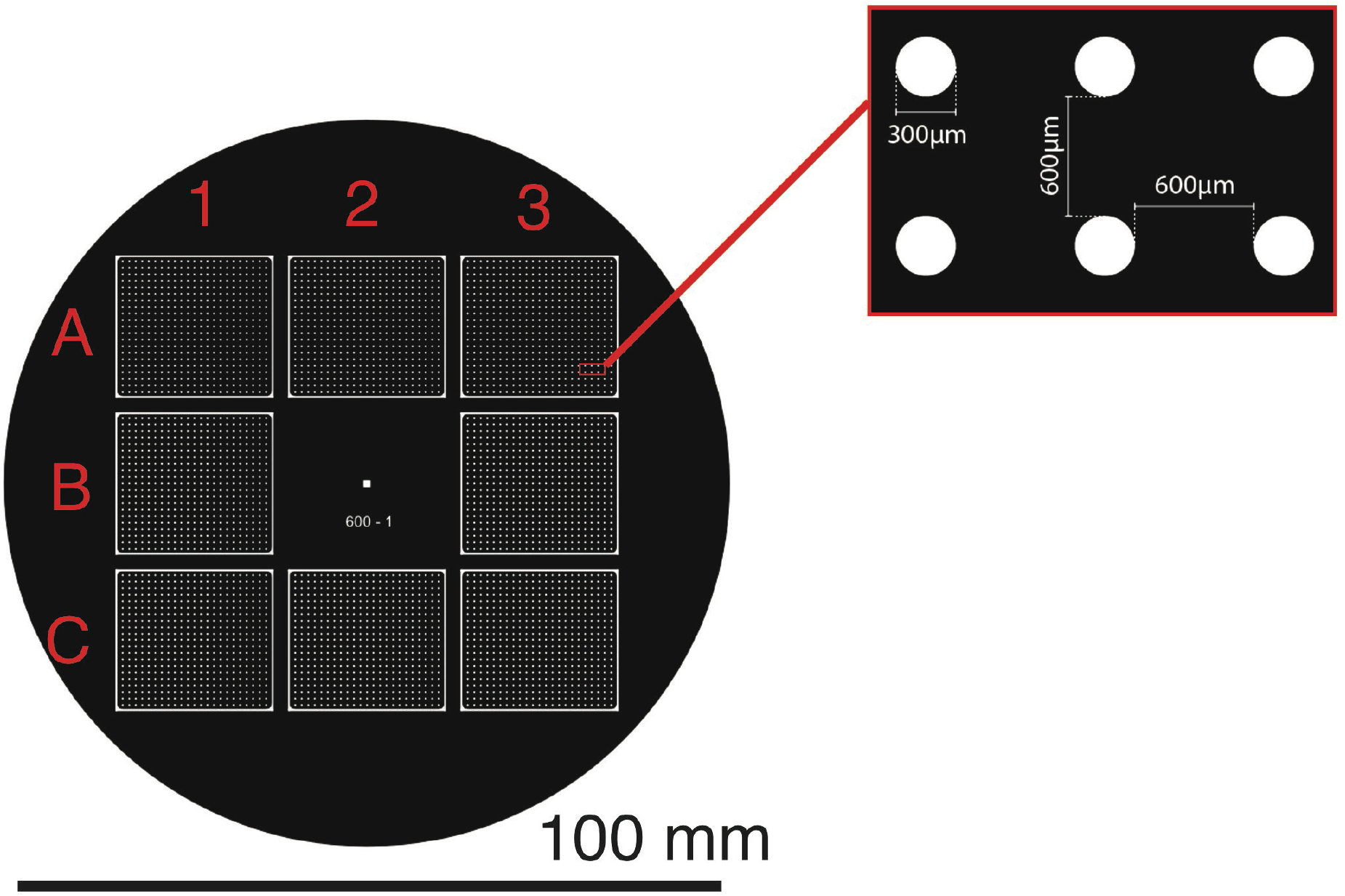
Photomask design. Diagram of the design of a photomask suitable for the round geometry of the silicon wafers (Siegert wafer) for 8 stencils with dimensions 21.3 × 19.1 mm each. The zoomed window shows the spacing of the micropattern with 300 μm holes separated by 600 μm spaces. Black areas represent opaque areas of the mask which match the shape of the wafer, while white areas allow UV exposure of the photoresist and will form the raised areas of the mould. Red letters and numbers are used to indicate the positions of the individual stencils in the grid. The photomask shown here is an example; the design should be tailored to the specific experiment.

- Check with your photomask manufacturer to ensure file type compatibility. We use LinkCad to convert files from dxf to Gerber format.
- The photomask design comprises a 3 × 3 array of rectangular stencils, each 21.3 × 19.1 mm (to fit the dimensions of 2-well ibidi slides) with a space in the centre (Figure 1). This design can produce eight stencils. Up to 4 of these designs can be ordered on a single 9 × 12” transparency.
- The photomask pattern for each stencil was designed such that there are 483 holes with a diameter of 300 μm each with a 600 μm spacing between the holes (Figure 1).
- The centre space was used for labelling but also contains a square of 1 × 1 mm in the centre, which will become a post for checking the height precision of the mould after manufacture.

## Key resources

| REAGENT or RESOURCE | SOURCE | IDENTIFIER |
| --- | --- | --- |
| **Chemicals** |  |  |
| SU8-GM1075 photoresist | Gersteltec | GM1075 |
| SU8 developer | Gersteltec | PGMEA |
| Isopropanol* | MicroChemicals GmbH | MIPU1025 |
| Trichloro(1H,1H,2H,2H-perfluorooctyl)-silane | Sigma Aldrich | 448931 |
| Sylgard 184 Silicone 1kg Elastomer Kit | Biesterfeld Spezialchemie | #G5498840000 |
| **Experimental models: Cell lines** |  |  |
| HM1 mouse embryonic stem cells |  | mouse ESC: HM1 |
| **Software and algorithms** |  |  |
| CAD software (e.g. CorelDraw, Adobe Illustrator, AutoCAD) | Corel/Adobe/AutoDesk | CAD software |
| **Other** |  |  |
| Film 9" x 12" Photomask * | JD Photo Data | Highest resolution 10 µm |
| Transparent glass plate * | JD Photodata | 125 mm x 125 mm x 2.35 mm |
| Silicon wafers 100 mm* | Siegert Wafer | BW14001 |
| White mixing cups 185 ml* | Hauschild | PP150-310 ml<br>#1000005111 |
| Square plastic petri dish* | Greiner | 688102 |
| Flat tweezers |  |  |
| Pointed tweezers |  |  |
| **Cell culture reagents** |  |  |
| µ-Slide 2 Well ibiTreat | ibidi | 80286 |
| 0.1% Gelatine in Water | Stem Cell Technologies | 07903 |
| DMEM/F-12, no glutamine | Thermo Fisher Scientific (Gibco) | 21331020 |
| Neurobasal (NB) Medium | Thermo Fisher Scientific (Gibco) | 21103049 |
| Accutase | Gibco | A1110501 |
| Bovine Serum Albumin (BSA) | Sigma-Aldrich | A3156 |
| Wash medium | 1:1 mix of DMEM/F12: NB medium, 0.1% BSA |  |
| N-2 Supplement (100X) | Gibco | 17502001 |
| B-27 Supplement (50X) | Gibco | 17504001 |
| GlutaMAX | Gibco | 35050061 |
| L-Glutamine | Gibco | 25030024 |
| Penicillin-Streptomycin (P-S) | Gibco | 15140122 |
| 2-Mercaptoethanol | Gibco | 31350010 |
| N2B27 medium | 1:1 mix DMEM/F-12:NB medium, 1X N2, 1X B27, 0.08% BSA, 2 mM L- |  |
|  | Glutamine or Glutamax, 100 U/ml P-S, 0.1 mM 2-Mercaptoethanol |  |
| bFGF | R&D | 3139–FB–025 |
| ROCK inhibitor Y-27632 | Tocris | 1254 |
| CHIR99021 | Axon | 1386 |
| 15ml falcon tubes* | Sarstedt | 62.554.502 |
| Cell Counting Slides* | Biorad | 1450011 |

\* = or equivalent

## Materials and equipment

| EQUIPMENT | SOURCE | IDENTIFIER |
| --- | --- | --- |
| Spin coater* | Polos | Spin150i |
| Digital programmable hotplate* | Harry Gestigkeit | 1°C resolution, 290 mm x 210 mm |
| EVG 610 Mask Aligner* | EVG, Austria | Mercury lamp |
| UV Filter* | Omega Optical | PL-360LP 215 mm x 215 mm |
| Crystallising Dish* | DURAN | 140 mm diameter #213135409<br>115 mm diameter #213134901 |
| Vacuum desiccator * | DURAN | NOVUS DN 200 clear |
| Nitrogen spray gun* | ipolymer.com Nitro-4 |  |
| Nitrogen spray gun * | Cleanroom world.com | with filter and Luer lock attachment TA-Nitro-3 |
| Benchtop Ultrasonic Cleaner* | Powersonic | P1100 with basket |
| Inspection microscope * | Nikon Eclipse L200N with 5x/10x objective |  |
| Microscope camera* | Optoteam | G5 |
| SpeedMixer* | Hauschild | DAC 150 |
| Desiccator* | Bel-Art | 230 mm Plate Size #999320237 |
| Spin Coater* | Laurell | WS-650-23 |
| Micrometer Screw Gauge | Mitutoyo | 1-micron resolution micrometer |
| Needle, blunted end* | McMaster Carr | 27G Luer lock |
| Convection incubator/oven reaching 80°C* | Binder | Model ED 56 |
| Inverted microscope* | Olympus CKX41 |  |
| Vacuum pump* |  |  |
| Chemical fume hood* | Wesemann |  |
| Benchtop centrifuge* | VWR | 521-1752 |
| TC20 automated cell counter* | Biorad | 1450102 |

\* = or equivalent

## Step-by-step method details

### Mould preparation (Timing: 1 day)

Fabrication of the mould uses photolithography. This procedure generates small, patterned structures from a light-sensitive material, photoresist (SU8). First, a silicon wafer is coated with a layer of photoresist, then baked. The photoresist is exposed to UV light through a photomask, which allows exposure in a specific pattern, and then baking again solidifies the pattern. Unexposed regions are dissolved in developer, leaving the desired pattern of the mould. The surface of the mould is coated with a hydrophobic layer to prevent unwanted adhesion of PDMS during stencil production.

#### Step 1: Prepare the photomask

##### NOTE

The photomask is a 0.18 mm thick polyester film with a photographic emulsion coated on one side.

1. Before use, cut out a single 4” (101.6 mm) photomask pattern from the film.
2. Tape the selected photomask flat onto the transparent 5” (127mm) glass plate with the printed surface facing away from the glass.

##### NOTE

The mask aligner chuck and mask tray can be used as a reference to align the mask on the glass plate so that it is perfectly aligned above the wafer in the mask aligner.

##### CRITICAL

Wear double gloves to avoid contamination of the developer and photoresist to your fingers.

#### Step 2: Dry silicon wafer

3. Set the hot plate to 110°C and bake the wafers for 5 mins to remove moisture and improve adhesion.
4. Cool the wafers to room temperature before use.

#### Step 3: Spin coat the wafer with SU8 photoresist

5. Centre the wafer on the chuck of the spin coater and apply a vacuum to hold it in place.
6. Dispense SU8-GM1075 onto the centre of the wafer by pouring from the bottle until approximately 40-50% of the diameter is covered.
7. Program the spin coater to spin at 500 rpm with an acceleration of 100 rpm/s for 110 s, followed by 900 rpm for 1 s with an acceleration of 400 rpm/s.
8. When the wafer is stationary, release the vacuum and move the wafer to a level surface at room temperature for a minimum of 10 mins until the photoresist is levelled.

##### NOTE

Air bubbles in the SU8 should be removed using a razor or needle to reduce defects in the final mould.

9. Transfer the wafer to the hot plate.
10. Ramp the temperature to 40°C over 5 mins.
11. Bake the wafer at 40°C for 30 mins.
12. Ramp the temperature to 120°C over 20 mins.
13. Bake the wafer at 120°C for 50 mins and then cool to room temperature.

##### NOTE

At this step, the mask aligner should be turned on to warm up the mercury lamp.

#### Step 4: Photolithography of pattern

In this step, a mask aligner is used to position the photomask over the SU8-coated wafer. UV light through the photomask patterns the SU8.

| Mask Aligner Setting | Value |
| --- | --- |
| Process | Man. Top Side |
| Process Mode | Transparent |
| Exposure Mode | Constant Dose |
| Contact Mode | Soft Contact |
| Mask holder | Size 5-inch |
| Chuck | Size 4 inch |
| Separation | 10 $\mu\text{m}$ |
| Thickness Mask | 2,3 mm |
| Thickness Substrate | 1 mm |
| Thickness Resist | 300 $\mu\text{m}$ |
| Exposure, Dose | 1500 mJ/cm<sup>2</sup> |
| Process | 600 mbar |
| WEC | 300 mbar |
| Exposure | 600 mbar |

14. Use the following settings for the mask aligner (EVG 610):
15. Follow the instructions on the mask aligner to load the glass plate with the photomask and SU8-coated wafer.
16. Add the UV filter (PL-360LP) to the light path.

##### NOTE

The UV filter prevents defects at the edges of patterns in the SU8 (T-topping).

##### NOTE

The soft contact mode on the mask aligner must be used to avoid distortion of the SU8.

17. Expose SU8 as defined in the mask aligner settings above.
18. After exposure, place the wafer on a hot plate at RT and set the temperature to 95°C. After the hot plate has warmed to 95°C, bake wafers for 2.5h.
19. Allow wafers to cool to room temperature.

#### Step 5: Develop photoresist to expose pattern

Following UV exposure and baking, the SU8 is ready for development. In combination with sonication, the developer dissolves SU8 that was not exposed to UV light.

20. Prepare 2 Crystallising Dishes of 115mm in diameter (small dishes) and one with a diameter of 140 mm (big dish).
21. Pour approximately 1 cm deep SU8 developer into the two smaller dishes. Pour approximately 1 cm height of isopropanol into the big dish.
22. Place one of the small dishes into the benchtop ultrasonic cleaner.
23. Place the mould in the dish in the ultrasonic cleaner.
24. Develop the SU8 for 16 mins at power 4 at RT.
25. Remove the mould from the ultrasonic cleaner and place it into the isopropanol dish to check for residues (brown-black stains fading into white).
26. Transfer the mould to the small dish with SU8 developer that was not in the benchtop cleaner to clean the residues by gently swirling the liquid around the wafer.
27. After all residues are dissolved, transfer the mould back to the isopropanol dish to check again.
28. If no white precipitate is forming, then wash once more in SU8 developer. Otherwise, repeat steps 26 and 27.
29. Remove the mould from the developer and dry the mould bottom with N_2_.
30. Mount the mould on the spin coater and spin at 3000 rpm for 30 s to dry.
31. Bake for 5 mins at 135°C on a hot plate to make SU8 permanent and avoid cracks in the material.
32. Check the mould under an inspection microscope for any irregularities in post diameters and inspect for damage.

#### Step 6: Silanization of the mould

Silanization of the mould adds a hydrophobic coating, which prevents the PDMS from sticking to the mould during stencil production.

##### NOTE

Trichloro-(1H,1H,2H,2H-perfluoroctyl)-silane is highly toxic to breathe or in contact with skin as it forms HCl with water. Use robust gloves and a fume hood.

33. In a fume hood, place the moulds in a desiccator connected to a vacuum pump.
34. Pipette 20 μl of Trichloro-(1H,1H,2H,2H-perfluoroctyl)-silane into a small tube and place it inside the desiccator without the lid.
35. Pull a vacuum of at least 100 mbar to vaporise the Trichloro-(1H,1H,2H,2H-perfluoroctyl)-silane.

##### NOTE

Reaching 100 mbar may take several mins, depending on the pump.

36. Seal the vacuum chamber.
37. After 1 h, let the air back in slowly.
38. To store the wafers, put them in separate square dishes lined with aluminium foil sheets at the bottom.

### PDMS Stencil Production (Timing: minimum 3h)

In this step, we use the coated wafers manufactured in the previous section to cast the PDMS stencils. Here, we spin-coat the mould with a layer of PDMS. PDMS that remains above the level of the posts on the mould must be removed by gentle blowing with N_2_ to ensure through holes in the final stencil. The PDMS is then baked to solidify.

1. In a white mixing cup, weigh the PDMS mixture (10:1 elastomer: curing agent) using Sylgard Elastomer 184 kit. Prepare approximately 10 g of PDMS per wafer.

#### NOTE

A plastic Pasteur pipette can be used to measure the volume of curing agent accurately.

2. Seal the lid and mix at 2000 rpm for 2 mins in a SpeedMixer.
3. Centre the mould on the chuck of the spin coater and apply vacuum.
4. Dispense the PDMS mixture onto the centre of the mould until approximately 50-70% of the diameter is covered.
5. Spin at 600 rpm for 30 s in a spin coater (for this, program the spin coater to 600 rpm for 40 s with an acceleration of 60 rpm/s).
6. Release the vacuum and remove the mould while avoiding PDMS drops. Place the mould in a square Petri dish on an aluminium foil sheet.
7. Apply a vacuum to the coated mould for 1 min to degas the PDMS.
8. Use a blunted 27G needle connected to an air gun at 10 psi to expose posts that PDMS covers. Hold the needle approximately 1 – 1.5 cm away from the mould. Blow over the mould in a regular grid pattern (up-down, left-right). Do two rounds on all moulds.

#### NOTE

The flow rate needs to be adjusted carefully to prevent the scattering of too much PDMS while ensuring exposure of the top of the pillars at the same time.

9. Bake the PDMS covered moulds in square dishes at 80°C for 2 h.
10. After cooling to room temperature, gently peel the stencils from a corner. Place the stencils flat onto a clean surface for storage.

#### NOTE

Moulds can be reused up to 6 times after careful cleaning of the residual PDMS.

11. The final PDMS stencils should be approximately 210 μm thick when measured with a micrometre screw gauge.

### Differentiation of spinal cord progenitors from mouse ES cells on stencils

The stencils can now be placed onto 2-well μ-slide ibiTreat dishes. The dish surface is coated overnight with gelatine and then dried to allow cell attachment. Stencils are placed with fine-tip tweezers.

#### Day 0: Coat dishes and sterilise stencils (Timing: 10 mins - 1 hour)

1. Coat the required number of 2-well μ-slide ibiTreat dishes with 0.1% Gelatine in H_2_O.

##### NOTE

Do not use gelatine in the PBS, as this will form crystals and prevent stencil attachment.

2. Leave in the incubator at 37°C overnight.
3. Sterilise selected stencils by UV in a laminar flow hood for minimum 20 mins to overnight.

#### Day 1: Plate stem cells onto stencils (Timing: 2-3h)

4. Aspirate the gelatine and leave dishes to dry at a 45° angle for 30-40 mins before placing stencils.
5. Carefully place the stencil using sterile forceps/tweezers.

##### NOTE

Ensure that the stencils are placed straight and without bulges and are not touching the edges of the dish, as this will lead to detachment.

##### NOTE

Due to surface tension, air will be trapped in the holes of the stencil.

6. To remove the air bubbles in the holes of the stencil, add 2 ml of a 1:1 mix of DMEM/F12 and Neural Basal media to the wells.

##### CRITICAL

Media **should not** contain BSA as this will act as a foaming agent.

7. Dishes (with lids off) are then placed into the desiccator inside sterile square plastic petri dishes.

##### NOTE

The square petri dishes allow the stacking of multiple dishes inside the desiccator.

8. Apply a vacuum for approximately 1 min.
9. Release vacuum. Most bubbles should be removed. Leave 1:1 media on the dishes and start preparing the ES cells.
10. Rinse cells with PBS.
11. Dissociate cells to a single-cell suspension using Accutase (1 ml per 10 cm dish).
12. Resuspend the cells in a total of 10 ml wash media and collect them in a 15 ml falcon tube.
13. Centrifuge cells at 1000 rpm for 4 mins using a dedicated cell culture centrifuge.
14. Aspirate supernatant and resuspend the cell pellet in 10 ml wash media.
15. Gently mix and load 10μl of the cell suspension into the counting slide chamber for automated counting. Count cells in duplicates.
16. Centrifuge cells again at 1000 rpm for 4 mins.
17. Resuspend cells at 1.25 - 1.5 million cells per ml in N2B27 medium supplemented with 10 ng/ml bFGF + 10μM Y-27632 ROCK inhibitor.
18. Aspirate the 1:1 medium from the ibidi dishes and add 2 ml of cell suspension per well. Let the cells settle for approximately 3 h in the incubator.
19. Wash cells twice with wash buffer to remove non-attached cells.

##### NOTE

Ensure that all floating cell clumps are removed, as those would reattach to your 2D colonies.

20. Add 2 ml N2B27 medium supplemented with 10 ng/ml bFGF and return cells to the incubator.

#### Day 2: Remove stencils (Timing: less than 30 mins)

21. Using sterile forceps, pinch one corner of the stencil and peel them off.

##### NOTE

Stencils are removed while still in media. Be careful not to disturb the attached colonies when peeling the stencil. Forceps with slightly bent tips work better for this as they naturally hook under the stencil.

22. Wash the cells with wash media.
23. Add N2B27 medium supplemented with 10 ng/ml bFGF and 5 μM CHIR and culture for 24h to obtain neuromesodermal progenitors.
24. After this point, continue with specific neural differentiation protocol. For full dorsal neural differentiation protocol, see Methods in ^9^.

## Anticipated results

Mould production should yield moulds with uniformly shaped and spaced pegs on top of the silicon wafer. Any defects in the mould will be transferred onto the stencils and subsequently affect the uniformity of the colonies.

Stencils production should result in PDMS stencils with uniform holes with clean well-defined edges (Fig. 2A). Good quality stencils will result in sharp-edged circular colonies that are restricted to the well boundaries and do not grow or spread under the stencil (Fig. 2B). Holes or clumps inside the patterned colonies results from suboptimal seeding densities.

**Figure 2.**
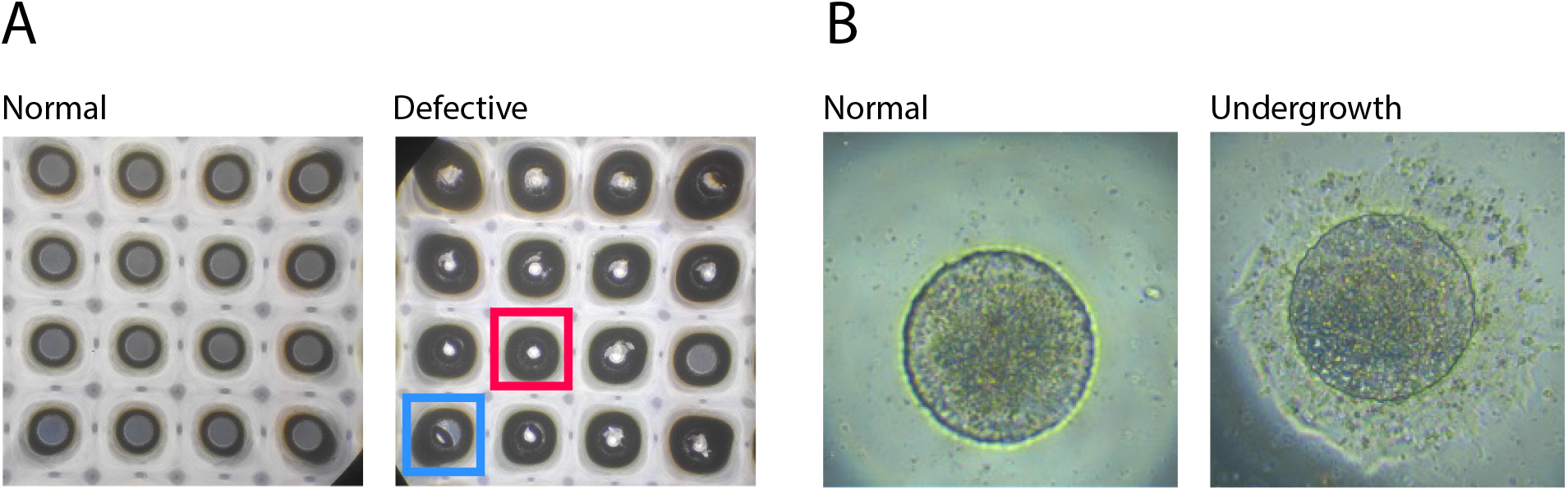
Potential stencil defects. **A) *Normal:*** Brightfield image of a good quality stencil with clearly defined holes. ***Defective:*** A bad stencil with multiple defects. The blue square shows a loose cap, and the red square shows a non-through hole, these present as a darker-rimmed hole with a light centre as they reflect the condenser of the microscope. **B) *Normal:*** Brightfield image of a colony before stencil removal on Day 2 (step 21). The colony is well contained within the well boundaries. ***Undergrowth:*** Brightfield image of a colony that has grown under the stencil, resulting in a poorly defined border. This results from poor stencil attachment to the dish.

Successful completion of the differentiation protocol should result in a high proportion of colonies with a radial pattern of expression of dorsal neural tube cell types. After stencil removal and subsequent 24 h treatment with RA and BMP4, as described in ^9^, neural crest cells are observed migrating outside of the colony, while roof plate cells are localised at the colony periphery. Within the colony, neural progenitor domains dP1 to dP6 form in their characteristic order^12^ from most dorsal at the periphery towards ventral in the colony centre (Fig. 3).

**Figure 3.**
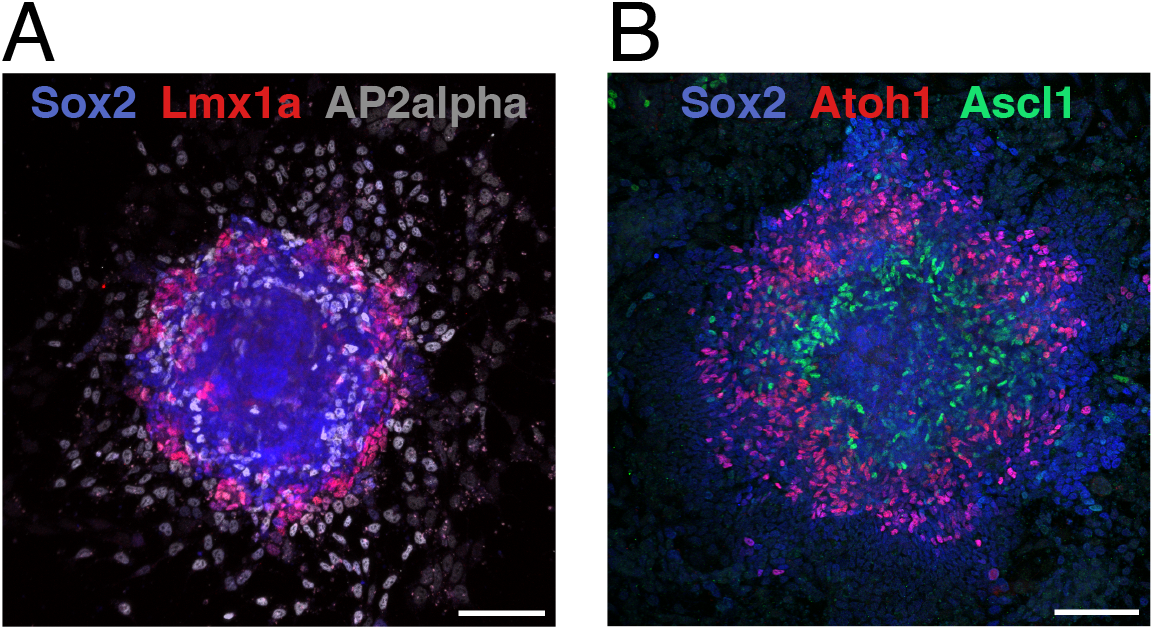
Self-organised patterns of neural crest and dorsal neural progenitor differentiation. Successful differentiation protocol results in self-organised radial patterns of dorsal neural tube cell types, such as the ones shown here. Immunofluorescence staining of dorsal neural tube progenitor domains. **A)** Colony at t = 48 h after the addition of 100nM RA and 0.5 ng/ml BMP4. Immunostaining against pan-neural progenitor marker Sox2, roof plate marker Lmx1a, and neural crest marker AP2alpha. **B)** Colony at t = 96 h after the addition of 100nM RA and 0.5 ng/ml BMP4. Immunostaining against pan-neural progenitor marker Sox2, dP1 marker Atoh1, and dP3-5 marker Ascl1. For antibody details as well as the specific details of neural differentiation, see ^9^. Scale bars, 100μm.

## Limitations

Using this method, it is challenging to achieve wells smaller than 300 μm as reduction of the post diameter or reducing any features to small dimensions produces a high height-to-width aspect ratio, which is more fragile.

The highest quality stencils are derived from the middle positions along the edges of the mould (Fig.1: A2, B1, B3 and C2), while corner stencils (Fig.1: A1, A3 and C1, C3) degrade in quality faster with repeated use. The centre position cannot be used for stencils due to limitations in spin coating efficacy in the centre of the wafer.

## Troubleshooting

Stencil defects usually arise at the blowing step (PDMS stencil production, step 8). Common defects are non-through holes (Fig. 2A, red square), which result from PDMS remaining on the top of the pillars. A related issue is caps (Fig. 2A, blue square); these can remain attached to the holes or become loose and stick to the stencils. Caps also result from poor clearing of the PDMS from the pillars but become dislodged during stencil demoulding.

Rough stencil edges can result from the underdevelopment of the mould or the bonding the PDMS to the SU8 due to poor salinisation.

Air bubbles in the PDMS next to the pillar will also ruin a stencil, but a vacuum treatment can remove them.

## Acknowledgements

We thank the nanofabrication facility at ISTA for technical assistance. Work in the AK lab is supported by ISTA, the European Research Council under Horizon Europe: grant 101044579, and Austrian Science Fund (FWF): Grant DOI 10.55776/F78. SL is supported by Gesellschaft für Forschungsförderung Niederösterreich m.b.H. fellowship SC19-011.

## Author contributions

SL and JM developed the method; SL, JM, MK, TM optimised the method; SL, JM, MK, TM and AK interpreted the results and wrote the manuscript; AK supervised the project and provided funding.

## Declaration of interests

No competing interests to disclose.

## Notes

### Competing Interest Statement

The authors have declared no competing interest.

